# Building the blood-brain barrier: a scalable self-assembling 3D model of the brain microvasculature under unidirectional flow

**DOI:** 10.1101/2025.09.10.675308

**Authors:** Jade Admiraal, Promise O. Emeh, Marleen Bokkers, Thomas Olivier, Karla Queiroz, Todd P. Burton, Nienke R. Wevers

**Author notes:** Authors contributed equally (Jade Admiraal, Promise O. Emeh, Marleen Bokkers).

## Abstract

The blood vessels of the central nervous (CNS) system form a tight, protective blood-brain barrier (BBB). This barrier is essential for healthy CNS function but also poses a hurdle in the treatment of increasingly common neurological disorders. Additionally, BBB dysfunction is a hallmark of many neurological diseases, further emphasizing a need for a better understanding of BBB function in health and disease.

We present a human self-assembling 3D model of the BBB in a microfluidic cell culture platform that allows culture of 48 models in parallel on one tissue culture plate. Human brain microvascular endothelial cells, pericytes, and astrocytes form highly reproducible BBB vascular networks under unidirectional perfusion and remain viable for a minimum of 14 days. Immunostaining reveals close cell-cell interactions with pericytes and astrocyte end-feet in direct contact with the brain microvasculature. Compared to endothelial monocultures, co-culture with astrocytes or pericytes results in improved barrier function, lower vessel diameters, increased branching, and alignment of the vessels in the direction of fluid flow. These results were most pronounced in tri-cultures containing all three cell types.

Unlike similar models previously reported, this brain microvasculature model allows for unidirectional perfusion without the need for pumps and syringes. Combined with its high-throughput nature, this feature renders the model suitable for studies of BBB function in health and disease, and assessment of potential BBB restorative therapies.

## INTRODUCTION

The cerebral blood vessels are formed by highly specialized endothelial cells. These cells are interconnected through tight and adherens junctions, creating a restrictive blood-brain barrier (BBB) (1,2). In addition to the endothelial layer, supporting cells such as pericytes and astrocytes play crucial roles in maintaining BBB integrity and function (3–5). The BBB protects the central nervous system (CNS) by preventing the uncontrolled entry of most circulating molecules and cells, thereby preserving brain homeostasis. Small lipophilic molecules can often cross the BBB via passive diffusion, whereas larger or polar compounds are mostly excluded. Essential molecules that cannot enter the brain via diffusion, enter via active transport mechanisms, such as glucose via the GLUT-1 transporter (4,6). In parallel, efflux transporters, such as P-glycoprotein and breast cancer resistance protein 1, actively expel various xenobiotics and drugs, further limiting CNS exposure (7,8).

While the restrictive nature of the BBB is essential for healthy brain function, it presents a major obstacle for the treatment of neurological diseases. Many therapeutic compounds administered systemically fail to reach the brain in concentrations high enough to elicit a therapeutic response (9,10). To address this challenge, various strategies have been developed to improve CNS drug delivery, but effective and predictable delivery across the BBB remains a major hurdle (11–13).

Disruption of BBB integrity is a hallmark of numerous neurological disorders, including Alzheimer’s disease, Parkinson’s disease, multiple sclerosis, and stroke (14,15). In these conditions, increased BBB permeability often coincides with neuroinflammation, exacerbating disease progression. Stroke ranks among the top three global causes of death and as a major cause of adult disability (16,17). At the same time, the number of deaths caused by dementias has doubled between 2009 and 2019 (18). The rising prevalence of such disorders underscores the urgent need for improved understanding of BBB function in both health and disease, as well as for the development of more effective CNS-targeted therapies.

Traditionally, much of our knowledge of the BBB has come from animal models. Although valuable, these models are expensive, time-consuming, and frequently limited by interspecies differences that reduce their translational relevance (19–21). To address these challenges, human *in vitro* BBB models have advanced considerably in recent years — evolving from simple monocultures to increasingly sophisticated co-cultures and microfluidic systems that more closely recapitulate the complexity of the *in vivo* BBB (22). Microfluidic models comprising endothelial cells, pericytes, and astrocytes show self-assembling BBB vascular networks that allow close cell-cell interactions and lumenized blood vessels (23,24). However, the lack of simple approaches for perfused cultured and low-throughput nature of these models poses a hurdle for broader adoption in the field, especially for compound-screening purposes.

Here we present a novel brain microvasculature model, using a microfluidic cell culture platform that enables the scalable generation of self-organized perfused vascular networks (25). This platform, called the OrganoPlate^®^ Graft 48 UF, harbors 48 chips in a single 384-well titer plate format and allows culture of 48 BBB vascular networks under unidirectional flow on one plate. Primary human microvascular brain endothelial cells (HBMECs) are combined with primary human pericytes and astrocytes and form highly reproducible vascular networks that remain viable for a minimum of 14 days. Dye perfusion shows retention of the dye within the brain microvasculature and bead perfusion reveals unidirectional flow through the vessels, mimicking cerebral blood flow. This model will allow studies of BBB development, its function in health and disease, and assess the potential of BBB restorative therapies.

## METHODS

### 2D Cell culture

Primary human brain microvascular endothelial cells (HBMEC, ACBRI 376, Cell Systems) were cultured in T75 flasks (F7552, Nunc^™^ Easy Flask, Sigma) in Promocell MV-2 medium (C-22121, Bioconnect). HBMEC were used for experiments between passage 7 and 8. T75 flasks (F7552, Nunc^™^ Easy Flask, Sigma) pre-coated with 50 µg/mL poly-L-lysine (3438-200-01, R&D Systems) were used for the expansion of primary human brain vascular pericytes (1200, Sciencell,) and human astrocytes (1800, Sciencell). Pericytes and astrocytes were cultured in their respective media (Pericyte Medium, 1201, Sciencell; Astrocyte Medium, 1801, Sciencell) and used between passage 4 and 5. All cells were maintained at 37°C, 5% CO_2_ and regularly tested for mycoplasma contamination and found negative. Cells were cultured for 4 days prior to seeding in the OrganoPlate^®^ and dissociated according to supplier’s instructions.

### OrganoPlate Graft 48 UF

The OrganoPlate^®^ Graft 48 UF (MIMETAS, Fig 1A, Suppl. Fig. 1) comprises 48 chips. Each chip consists of a gel chamber in which cells embedded in extracellular matrix gel can be added, and two adjacent perfusion lanes. Phaseguides positioned between the gel chamber and perfusion lanes function as capillary pressure barriers to pattern the gel and prevent it from flowing into the adjacent perfusion lanes (26). Each chip comprises one longer and one shorter perfusion lane that both have inlets and outlets in long wells (3 wells of a 384-well plate merged together). The long wells (Fig. 1C, wells 1-3 and wells 6-8) combined with a low media volume allow for creating an air–liquid interface above the outermost holes in the chip at positions 1 or 8 when the plate is angled above a critical angle. The formation of an air–liquid interface above an inlet hole prevents fluid flow, and the size constraints of the device result in a Laplace value where surface tension prevents the microfluidic channel from emptying. When the angle of inclination is negative, fluid is routed through the short channel and split down the two paths: (1) through the gel chamber and towards the long channel and (2) continuing through the short channel. The mode of action is mirrored in the positive incline (Fig. 1D).

**Figure 1.**
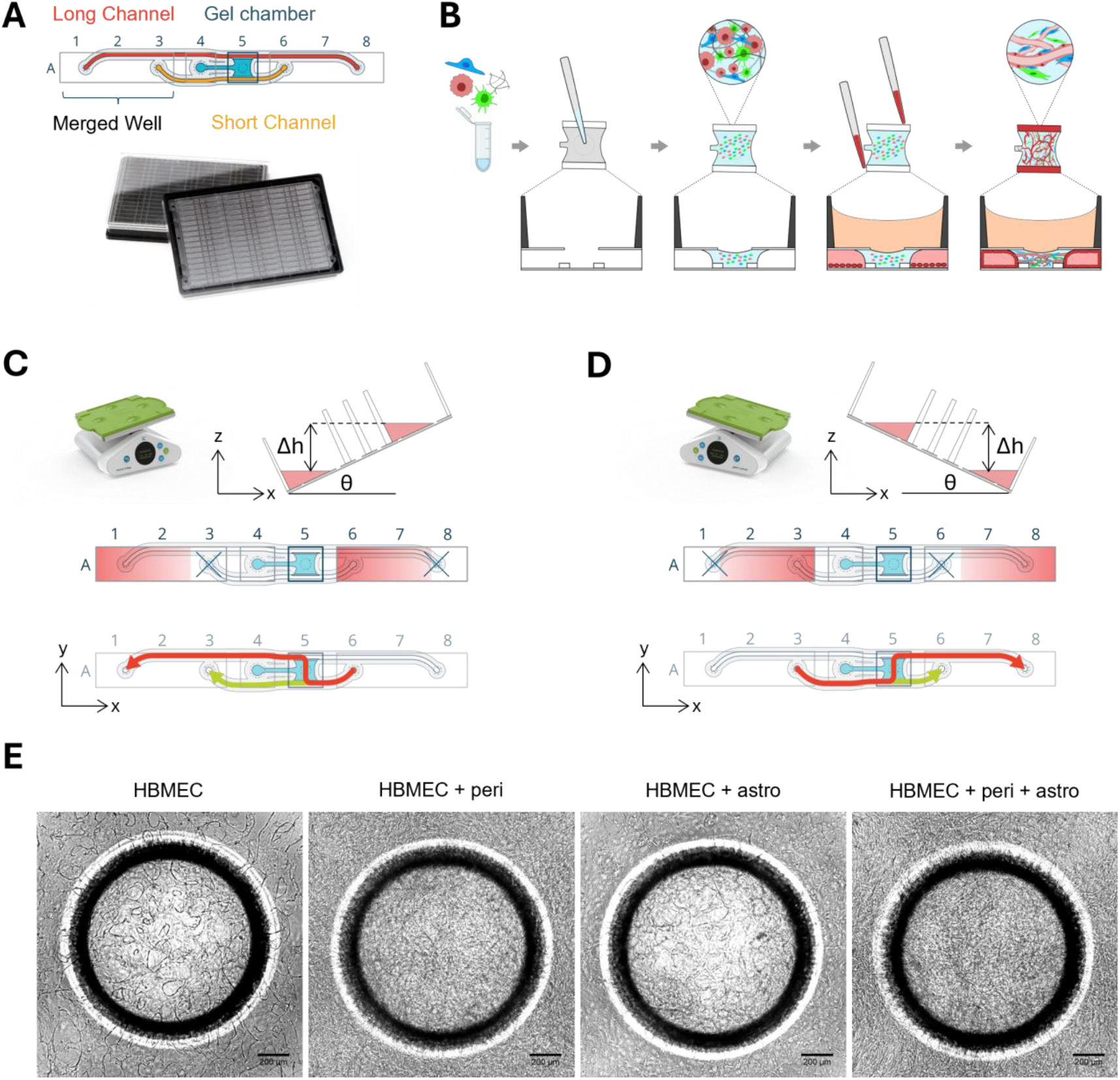
Brain-derived cells self-assemble into vascular networks in the OrganoPlate Graft 48 Uniflow. **(A)** The OrganoPlate Graft 48 UF is based on a modified 384-well microtiter plate format where long wells are created by merging together 3 wells of the titer plate to link perfusion channels fluidically. It contains 48 chips that can be used to model a BBB vascular network. Each chip comprises a gel chamber (blue) and two perfusion lanes (red, orange). **(B)** Human brain microvascular endothelial cells (HBMECs), astrocytes, and pericytes are embedded in an extracellular matrix gel and seeded to the gel chamber of each chip. Next, HBMECs are seeded in the adjacent perfusion lanes and form vessel-like structures against the extracellular matrix gel. The cells self-assemble into a 3D BBB network. **(C)** Perfusion is generated by placing the OrganoPlate Graft 48 UF on a rocker platform that induces unidirectional gravity-driven flow through the BBB microvasculature in the gel chamber. During the negative incline, flow is driven from the right long well towards the left long well, with the rightmost hole at A8 becoming disconnected from the fluid in the well, creating a pressure drop across the gel chamber. **(D)** In the positive incline, flow is driven from the left long well towards the right long well and a pressure drop is created across the gel chamber when the hole at A1 is disconnected from the fluid in the well due to the inclination angle of the microtiter plate. **(E)** Phase contrast pictures of BBB monocultures and co-cultures obtained on day 7 after seeding. Scale bar = 200 µm.

### OrganoPlate culture

HMBECs, pericytes, and astrocytes were dissociated from flasks and embedded in a mixture of fibrinogen (F3879, Sigma-Aldrich), thrombin (HT1002a, Enzyme Research Laboratories), and Matrigel-GFR (356231, Corning). The resulting gel had a final concentration of 5 mg/mL fibrinogen, 0.2 U/mL thrombin, and 5% (v/v) Matrigel-GFR. HBMECs were seeded at 10,000 cells/µL, pericytes at 2,500 cells/µL, and astrocytes at 500 cells/µL throughout all tested conditions (HBMEC monoculture; HBMEC-pericyte coculture; HBMEC-astrocyte coculture; and HBMEC-pericyte-astrocyte co-culture). 1.5 µL cell-ECM mixture was loaded into the gel chamber of the OrganoPlate^®^ Graft 48 UF (4801400B, MIMETAS). Plates were incubated for 15 minutes in a humidified incubator (37°C, 5% CO_2_), after which 50 µL of MV-2 medium was added to the gel chamber. A Matrigel-GFR solution (1:100 in PBS, 70013065, Gibco) was then added to the perfusion channels for coating and incubated for 1 hour. The coating solution was removed and HBMECs were seeded in the perfusion channels at a density of 10,000 cells/µl using passive pumping technique (27). Plates were then incubated for 2-3 hours (37°C, 5% CO_2_) to allow HBMEC attachment. Next, 100 µL Promocell ECGM-2 medium (C-22011, Promocell) was added to the in- and outlets of both perfusion channels. The medium in the gel chamber was replaced with 50 µL ECGM2 supplemented with 1% pericyte growth supplement (1252, ScienCell) and the plate was placed on an OrganoFlow^®^ perfusion rocker (MI-OFPR-L, MIMETAS) set at 7° inclination, 8-min interval to media perfusion (bi-directional for first 24h). One day post seeding, the medium volumes were changed to 40 µL in the perfusion channels and 20 µl in the gel chamber and the plates were placed on an OrganoFlow rocker set at 25° inclination, 1-min interval to allow for unidirectional perfusion to the gel chamber. From day 1-7, the ECGM2 medium added to the perfusion lanes was supplemented with 50 ng/mL VEGF (100-20, PeproTech), 250 nM S1P (73914, Sigma-Aldrich) and 100 kIU/mL aprotinin (7005124, Nordic Pharma), while for the gel chamber the ECGM2 was supplemented with aprotinin. After day 7, VEGF and S1P were removed from the medium. Medium changes were performed thrice weekly.

### Immunocytochemistry

Cultures were fixed using 3.7% formaldehyde (252549, Sigma-Aldrich) or 100% methanol (-20°C, 494437, Sigma-Aldrich) and incubated for 15 min. Cultures were permeabilized and blocked for 2 hours by adding PBS containing 1% Triton X-100 (T8787, Sigma Aldrich) and 3% bovine serum albumin (BSA, A2153, Sigma-Aldrich). Primary antibodies were prepared in an antibody incubation buffer consisting of 0.3% Triton X-100 and 3% BSA in PBS and incubated at RT overnight. The following primary antibodies were used: mouse anti-PECAM-1 (M0823, DAKO), rabbit anti-VE-cadherin (Ab33168, Abcam), mouse anti-Claudin-5 (35-2500, Thermo Fisher Scientific), rabbit anti-PDGFRβ (Ab32570, Abcam), chicken anti-GFAP (Ab4674, Abcam) and rabbit anti-AQP4 (PA5-53234, Invitrogen). Next, chips were washed 3x with PBS supplemented with 0.3% Triton X-100 (washing solution) and secondary antibodies were applied in antibody incubation buffer and incubated together with NucBlue^™^ reagent (R37606, Thermo Fisher Scientific) overnight at RT. The following secondary antibodies were used: donkey anti-mouse 488 (A21202, Thermo Fisher Scientific), donkey anti-rabbit 750 (ab175731, Abcam), goat anti-chicken 647 (RA21449, Thermo Fisher Scientific), ActinRed^™^ 555 reagent (R37112, Thermo Fisher Scientific). Finally, chips were washed with washing solution followed by PBS, after which plates were imaged using the Confocal Micro XLS-C high content imaging systems (Molecular Devices, z-step height 5 µm).

### Immunostaining quantification

PECAM-1 images obtained from mono- and co-cultures were quantified using Fiji (28). The images were pre-processed to reduce background signal by applying a rolling ball background correction routine (29). The center of the gel chamber was selected and segmented using a trainable classifier in Labkit (30,31). Vessel signal was extracted as foreground (white), and non-vessel signal was considered background (black). The segmented vessel signal was then skeletonized, after which vascular characteristics were extracted. Extracted values were then compared across BBB cultures and were plotted using GraphPad Prism, version 10.

### Dye perfusion assay

To confirm the presence of perfusable vessels, 40 µL of ECGM-2 medium containing 0.25 mg/mL FITC-Dextran (150kDa, 46946, Sigma-Aldrich) was added to the perfusion inlets and outlets, while the gel chambers remained empty. The plates were then placed on the OrganoFlow rocker (25° inclination, 1-min interval) for 4 minutes to enable dye perfusion through the established vascular network. Fluorescent images were acquired using an ImageXpress Micro XLS-C system (Molecular Devices) at 37°C with a 4X objective.

### Bead perfusion assay

To confirm the presence of perfusable vessels, 40 µL of ECGM2-medium containing 1-5 µm yellow fluorescent polymer microspheres (FMY, Cospheric) was added to perfusion channels inlets and outlets, while the gel chambers remained empty. The plates were then placed on a rocker (set to 25 degrees, 1 min) to enable perfusion through the established vascular network and were monitored using a EVOS^™^ FL Auto 2 (AMAFD2000, Thermo Fisher Scientific) with a 4X objective. Recordings were captured at 20 frames per second for each of the culture conditions.

The recordings were pre-processed by using a Kalman filter (32) to reduce high gain noise from the time lapse recording and used for bead perfusion quantification. The angular local orientations of the bead flow were then quantified using OrientationJ (33–35). The visual directional analysis was used to create example images of each culture condition, overlaying local orientations as colors (hue) on the original image. The vector field implementation was used to quantify the local orientations and was measured for each frame of the recording, and post-processed using Python.

### Statistical analysis

Data was analyzed using GraphPad Prism, version 10. Gaussian distribution was assessed using the Shapiro-Wilk normality test. A Brown-Forsythe and Welch test including a Dunnett T3 multiple comparisons test was performed in case of normally distributed data in which the assumption of equality of variances was violated (figure 2d-f). When normal distribution could not be confirmed (figure 2c,), the nonparametric Kruskal-Wallis test with Dunn’s multiple comparisons test was performed. Statistical significance was indicated by one or more asterisks. *(P < 0.05), **(P < 0.01), ***(P < 0.001), or ****(P <0.0001).

**Figure 2.**
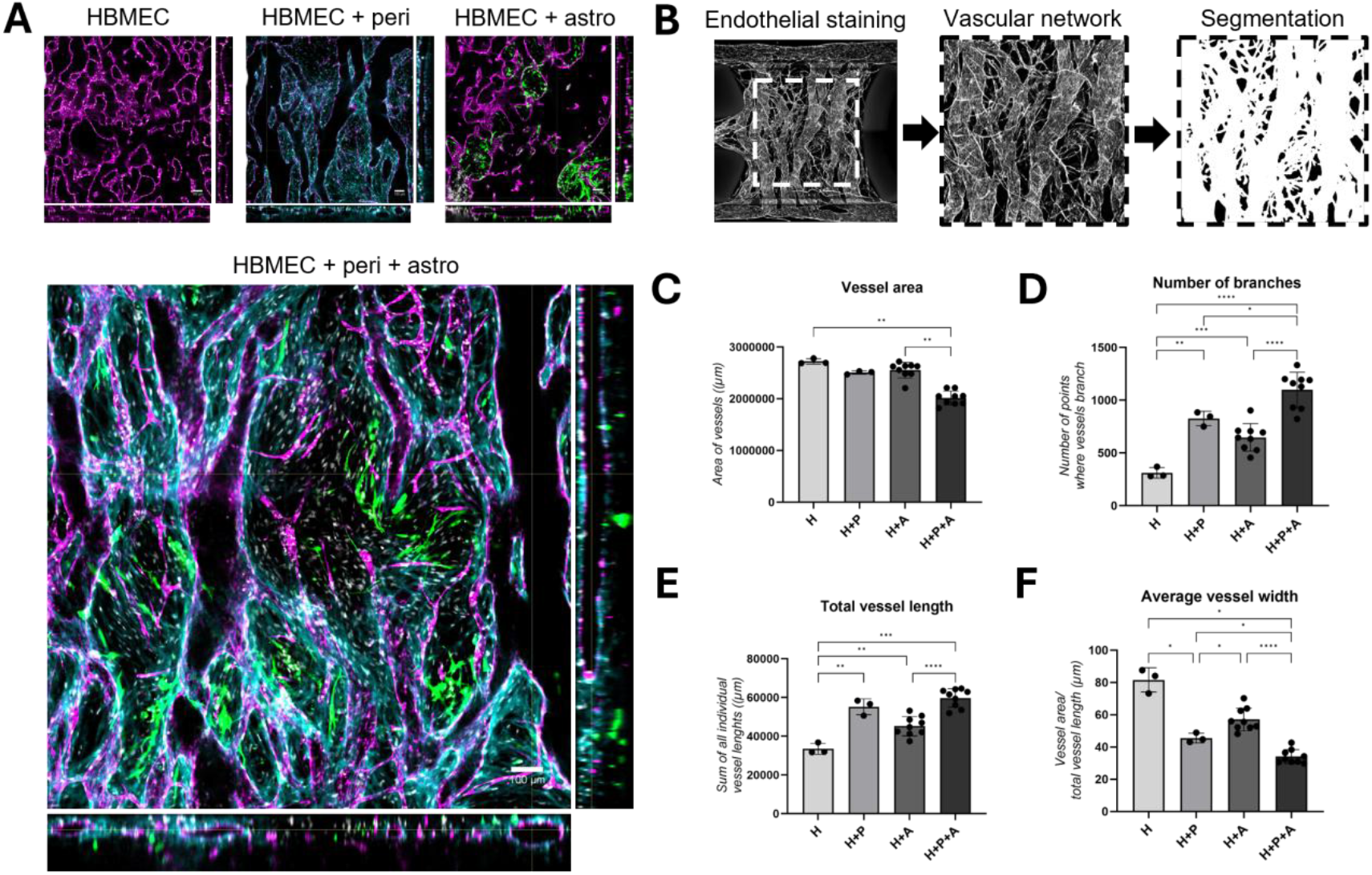
Vascular networks show brain endothelial vessels in close contact with pericytes and astrocytes. **(A)** Cerebral vascular networks consisting of (1) HBMECs, (2) HBMECs + pericytes, (3) HBMEC +astrocytes, or (4) HBMECs + pericytes + astrocytes) were cultured for 14 days and stained for endothelial cell marker PECAM-1 (magenta), pericytic marker PDGFRβ (cyan), and astrocytic marker GFAP (green). Orthogonal projections display single-slice images of the top view, front view, and side views of the gel chamber. Scale bar = 100 µm. **(B)** Maximum projection images of the vasculature (PECAM-1) were used for quantification purposes following a segmentation procedure. **(C)** Quantification of vascular network area. **(D)** Quantification of total number of branches. **(E)** Quantification of total vessel length. **(F)** Quantification of average vessel width of different BBB cultures. Data represents the mean ± SD for n=3-9 chips, *(P < 0.05), **(P < 0.01), ***(P < 0.001), ***(P < 0.0001).

## RESULTS

### Brain endothelial cells, pericytes, and astrocytes self-assemble into vascular networks in the OrganoPlate Graft Uniflow

We employed a novel 3D cell culture system, called the OrganoPlate Graft Uniflow (UF). This plate harbors 48 microfluidic chips in a 384-well microtiter plate format (Fig. 1A, Suppl. Fig. 1). Primary human brain microvascular endothelial cells (HBMECs), pericytes, and astrocytes were embedded in an extracellular matrix (ECM) gel consisting of fibrin and Matrigel and loaded into the gel chamber of each chip. After gelation, HBMECs are added to the two adjacent perfusion lanes (Fig. 1B). Perfusion of the cultures is initiated by placing the OrganoPlate Graft UF on a gravity-driven perfusion rocker that induces bidirectional flow through the perfusion lanes harboring HBMECs, but unidirectional flow through the gel chamber depending on the positive or negative rocking angle (Fig1. C-D). This results in the formation of an unidirectionally-perfused ECM-embedded BBB vascular network, connected to two larger endothelial vessels (Fig. 1E).

### Vascular networks show brain endothelial vessels in close contact with pericytes and astrocytes

Cerebral networks consisting of (1) HBMECs only, (2) HBMECs and pericytes, (3) HBMEC and astrocytes, or (4) HBMECS, pericytes, and astrocytes were cultured in the OrganoPlate Graft UF for 14 days and characterized using immunofluorescent staining. Confocal imaging revealed hollow structures lined by PECAM-1 positive cells, indicating a network of lumenized endothelial vessels, in all tested culture setups (Fig. 2A). In addition, the vessels showed expression of endothelial marker VE-cadherin as well as tight junction protein Claudin-5 (Suppl. Fig. 2A), confirming a BBB-specific endothelial phenotype. Pericytic marker PDGFRβ is observed on the outside of the endothelial vessels, directly in contact with the endothelial cells, as are the end-feet of astrocytes, expressing GFAP and AQ4 (Fig. 2A, Suppl. Fig. 2A). To further characterize the vascular networks, PECAM-1 staining images were converted into binary images in which white signal indicated the presence of vessels with their branches (Fig. 2B, Suppl. Fig 2B). A significant reduction in vascular network area was observed with the addition of pericytes and astrocytes compared to HBMEC monocultures (P= 0.0017 for triculture versus monoculture) (Fig. 2C). Additionally, the brain microvasculature of BBB co-cultures showed a significant increase in vascular branching compared to HBMEC monocultures, especially in the tricultures (1098 branching points vs. 312 branching points, P≤0.0001) (Fig. 2D). Finally, addition of pericytes and astrocytes resulted in a significant increase in total vessel length (P=0.0001) and a reduction in average vessel width compared to HBMEC monocultures (34 µm in tri-cultures vs. 82 µm in monocultures, P=0.0272) (Fig. 2F). Together, these data show more intricate microvascular network formation in presence of astrocytes and pericytes.

### Robust formation and perfusion of cerebral vascular networks

To confirm that the hollow vessel structures as observed via immunostaining were functional, the cerebral vascular networks were perfused with a fluorescent dye. A minimum of 20 chips were assessed per culture condition at 4 different time points after cell seeding. Perfusion of the BBB vascular networks showed retention of the dye in the vessels’ lumens with high reproducibility (Fig. 3A). While HBMEC monocultures and co-cultures of HBMECs and astrocytes showed moderate leakage of the dye from the vascular networks into the ECM gel, co-cultures containing pericytes retained the dye in their lumens. All BBB vascular networks models showed robust perfusion (>95% chips) over the assessed period of day 7-14 (Fig. 3B). HBMEC monocultures and co-cultures of HBMECs with astrocytes show slightly reduced leak tightness over time. In contrast, vascular networks containing pericytes remain leak-tight, showing no leakage of dye into the ECM gel area at any of the tested timepoints. Additionally, pericyte-containing co-cultures show vascular remodeling and pruning of the network as well as contraction of the two larger vessels in the perfusion lanes, likely due to the contractile nature of the pericytes.

**Figure 3.**
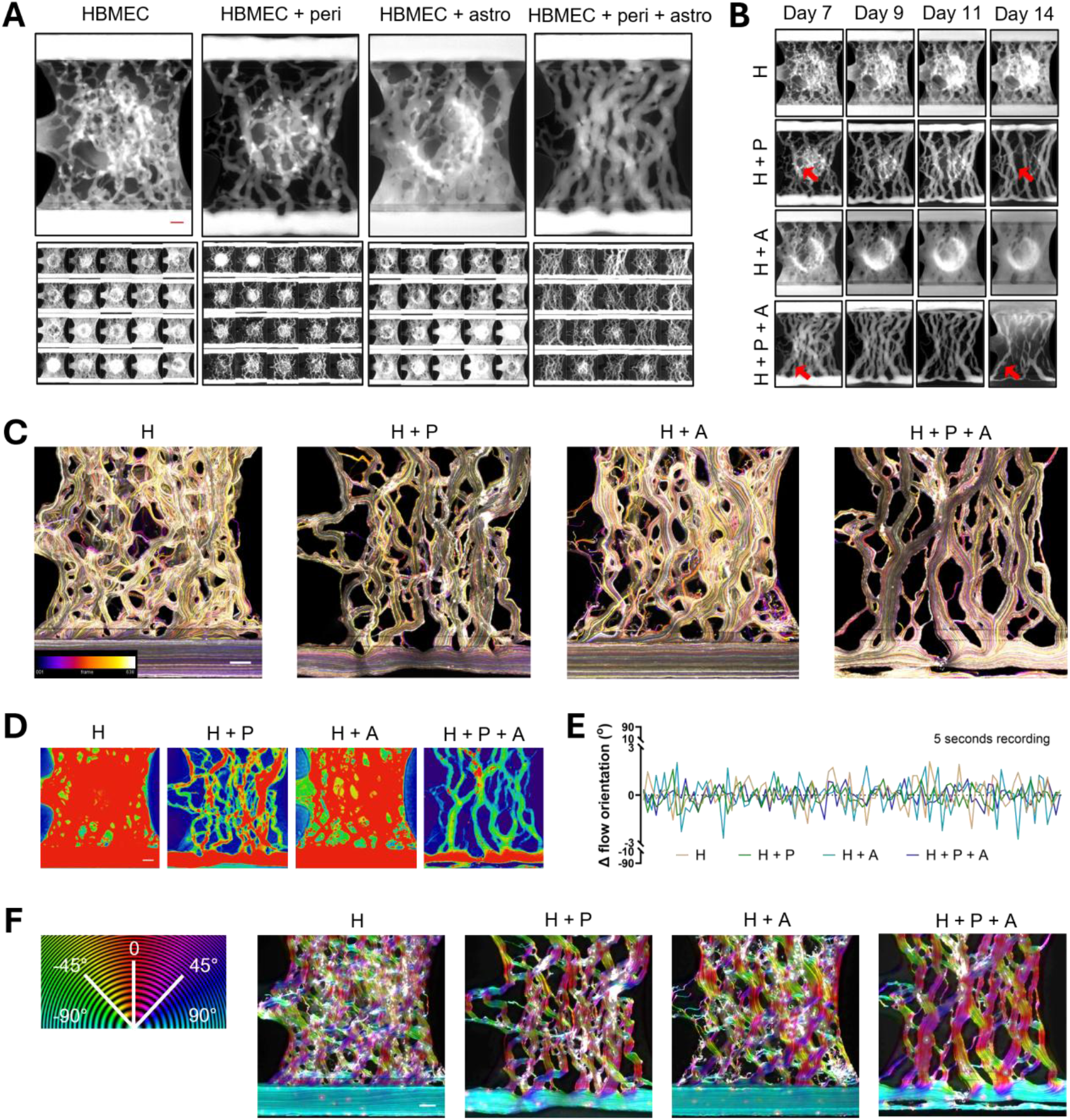
Robust brain microvasculature formation and perfusion. **(A)** 150 kDa FITC-dextran dye was perfused through BBB vascular networks consisting of (1) HBMECs, (2) HBMECs + pericytes, (3) HBMECs + astrocytes, and (4) HBMECs + pericytes + astrocytes. Upper panels show representative images for each culture condition obtained 4 min after dye addition. Lower panels show a montage of 20 assessed chips for each condition at day 7 of culture. **(B)** Perfusion of the different BBB vascular networks with a 150 kDa FITC-dextran dye was assessed at different time points (day 7-14). Arrows indicate areas that show vascular remodeling and pruning over time. **(C)** Fluorescent beads were perfused through the cerebral vascular networks. Bead flow was captured via high-speed fluorescent imaging. Colors correspond with frame number in the color-coded image indicated in the colormap. **(D)** Heatmaps show low-traffic (blue) and high-traffic (red) bead flow areas in BBB vascular networks consisting of (1) HBMECs, (2) HBMECs + pericytes, (3) HBMECs + astrocytes, and (4) HBMECs + pericytes + astrocytes. **(E)** Change in flow orientation over time. A value close to 0 means limited directional changes are found relative to the previous frame, and values deviating from 0 mean significant angular changes compared to the previous frame. **(F)** Flow directions were color coded for each culture setup, with flow from right to left (-90°) and left to right (90°) represented in teal and flow straight through the gel chamber (0°) depicted in red. White colors imply no direction can be determined for that location. Scale bar = 200 µm.

In addition, flow direction through the vascular networks was assessed via perfusion with fluorescent microsphere beads of 1-5 microns in size. Beads were added to the perfusion lanes and their flow through the chips was monitored via high-speed fluorescent imaging. The fluorescent beads reveal a unidirectional flow through the BBB vascular networks, entering from the bottom perfusion lane and exiting via the top perfusion lane (Suppl. Videos 1-4, Fig. 3C). High- and low-bead traffic areas were visualized using a heatmap approach (Fig. 3D) and show higher bead traffic in networks devoid of pericytes, in line with the higher vessel width and lower vascular branching observed in these conditions (Fig. 2D&F). Quantification (Fig. 3E) and color-coded visualization (Fig. 3F) of bead flow confirmed that the overall flow patterns through the vascular networks are unidirectional in nature for all BBB models. More detailed assessment revealed that beads perfused through HBMEC monocultures and co-culture of HBMECs and astrocytes showed more variation in flow direction compared to those perfused through cultures containing pericytes, which show alignment of the vessels in the direction of fluid flow. This result was most pronounced in tri-cultures containing all three cell types.

## DISCUSSION

In this study, we present a 3D self-assembling blood–brain barrier (BBB) model comprising primary human brain endothelial cells, pericytes, and astrocytes cultured under gravity-driven unidirectional flow. Immunostaining revealed a network of endothelial vessels in direct contact with pericytes and astrocyte end-feet, embedded in ECM gel. The endothelial networks consist of functional vessels as evidenced by the perfusion with fluorescent dye and beads. Using a novel chip design, we show unidirectional flow through the BBB networks without the need for pumps or syringes.

A key focus of this work was to investigate the influence of pericytes and astrocytes – separately and in combination – on the formation, morphology, and functional integrity of the BBB vascular network. All tested culture configurations, including (1) HBMEC monocultures, (2) HBMEC + pericyte co-cultures, (3) HBMEC + astrocyte co-cultures, and (4) HBMEC + pericyte + astrocyte co-cultures robustly formed perfusable cerebral networks. Co-culture conditions showed stratified organization of pericytes and astrocytes surrounding the endothelial vessels with direct contact verified through immunocytochemistry characterization. This structural mimicry is essential, as both cell types are known to regulate BBB integrity, basement membrane integrity, and BBB-specific gene expression through paracrine signaling and physical contact (36–38). HBMEC monocultures and co-cultures of HBMECs and astrocytes presented with lower barrier integrity, larger vessel diameters, and fewer branches compared to co-cultures containing pericytes. Addition of pericytes induced tighter barrier function, smaller vessel diameters, and more branching. These effects were most pronounced in tricultures containing all three cell types. Together, these data show more intricate microvascular network formation in presence of astrocytes and pericytes. Addition of pericytes induced tighter barrier function, smaller vessel diameters, and more branching. These effects were most pronounced in tricultures containing all three cell types. Together, these data show more intricate microvascular network formation in presence of astrocytes and pericytes.

Furthermore, the inclusion of pericytes – in absence or presence of astrocytes – resulted in vascular structures that were more aligned in the direction of flow. This observation suggests that pericytes may play a critical role in guiding endothelial remodeling and axial alignment in response to mechanical cues, consistent with *in vivo* studies highlighting pericyte involvement in vascular development, maturation and stabilization (39,40). The observed synergistic effects may also suggest a pivotal role of astrocytes in vascular organization by ensheathing blood vessels and regulating cerebral flow although their function is context-dependent as it is influenced by developmental-maturational stage, interactions with other cell types and pathological conditions (41–43). One must note that the cell ratios used in this study were selected based on *in vitro* optimization and were not equal for astrocytes (500 cells/µL) and pericytes (2,500 cells/µL), complicating direct comparison of HBMEC + astrocyte and HBMEC + pericyte co-cultures.

Several other reports previously described the formation of self-assembling BBB vascular networks in fibrin-based ECM gels (23,24,44). However, these made use of polydimethylsiloxane (PDMS) based chips, which suffer from hydrophobicity and fouling problems, are low in throughput, and require complex setups to generate perfusion flow. Very recently, Rajput et al. described a self-assembling BBB vascular network model in the OrganoPlate^®^ Graft using brain endothelial cells and pericytes to study the effect of Flavivirus on BBB dysfunction (45). Unlike most microfluidic chips, this system allows higher throughput experimentation and makes use of gravity-driven medium perfusion.

The model described in the present manuscript shows high resemblance to the recent work of Rajput et al. but utilizes a novel chip design, the OrganoPlate^®^ Graft Uniflow, that enables a unidirectional rather than bidirectional fluid flow through the microvasculature. The unidirectional flow better mimics the cerebral blood flow *in vivo* and may allow for improved assessment of vascular function, as flow disturbances have been shown to induce vascular dysfunction *in vivo (46)*. Our simple yet physiologically relevant setup supports continuous perfusion through the lumenized vascular network, making it cost-effective, scalable, and well-suited for studying immune cell trafficking and drug permeability studies under dynamic flow conditions. Furthermore, our work incorporates astrocytes alongside pericytes and endothelial cells, enabling further research into pericytes’ and astrocytes’ distinct and synergistic contributions to BBB function in health and disease.

The model presented here may be applied to study common neurological diseases, such as stroke, using an *in vitro* approach that mimics reduced perfusion and oxygen deprivation (47). Additionally, the gravity-based perfusion flow is well suited for assessing adhesion and extravasation of circulating immune cells such as leukocytes into the brain compartment (48). While the current cerebral vasculature model incorporates several key cellular structures of the human BBB, the inclusion of additional cell types involved in the neurovascular unit (NVU), such as neurons and microglia, would further improve the physiological relevance of the model (49–51).

## CONCLUSION

We present a 14-day stable, highly robust self-assembling BBB co-culture model containing primary human brain endothelial cells, pericytes, and astrocytes. This model allows for assessment of each cell type’s distinct contribution as well as their synergistic effects. With 48 chips in one plate and gravity driven unidirectional flow through the cerebral vascular network, this model poses a scalable and applicable tool in the study of BBB dysfunction and restorative therapies.

## LIST OF ABBREVIATIONS

AQ4: aquaporin 4
BBB: blood-brain barrier
CNS: central nervous system
ECM: extracellular matrix gel
HBMEC: human brain microvascular endothelial cell
PECAM-1: platelet endothelial cell adhesion molecule
PDGFRβ: platelet derived growth factor receptor beta
GFAP: glial fibrillary acidic protein
PDMS: Polydimethylsiloxane
VE-cadherin: vascular endothelial cadherin
UF: uniflow.

## DATA AVAILABILITY

The datasets used and/or analyzed during the current study are available from the corresponding author on reasonable request.

## DECLARATIONS

### Ethics approval and consent to participate

Not applicable.

### Consent for publication

Not applicable

### Author contributions

Authors J.A., P.O.E., and M.B. performed experiments and data analysis and therefore share first authorship. T.O. performed image analysis and quantification. T.P.B. designed the microfluidic chip used in this study. K.Q., T.P.B., and N.R.W. designed and supervised the research. All authors wrote and approved the final manuscript.

## Acknowledgements

The authors thank Artur Rodrigues for his help with bead perfusion assays, Alexis Dalaud, Kesley Bouwmeister and Thijs Slot for chip fabrication, Dr. Xandor Spijkers for the experimental support and Prof. Elga de Vries, Dr. Martine Lamfers and Prof. Clemens Dirven for their valuable input and discussion of this work.

