## Supplementary figure 1 and 2 for "Building the blood-brain barrier: a scalable self-assembling 3D model of the brain microvasculature under unidirectional flow"

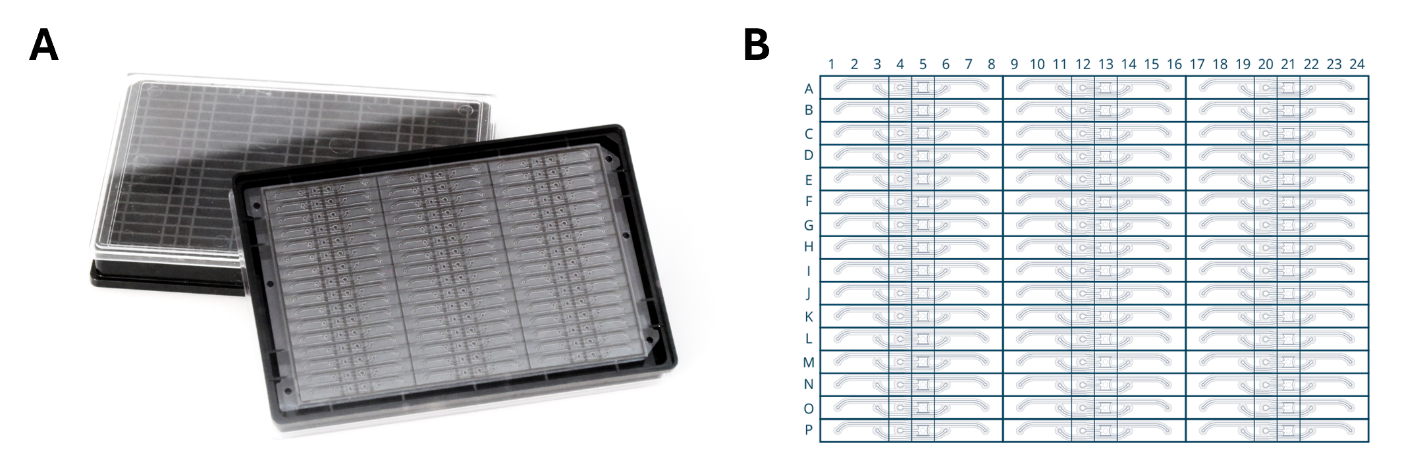

**Supplementary Figure 1 | The OrganoPlate Graft 48 UF.** **(A)** Top and bottom view of the OrganoPlate Graft 48 UF. **(B)** Plate layout of the OrganoPlate Graft 48 UF showing 48 individual tissue culture chips in the format of a modified 384-well culture plate.

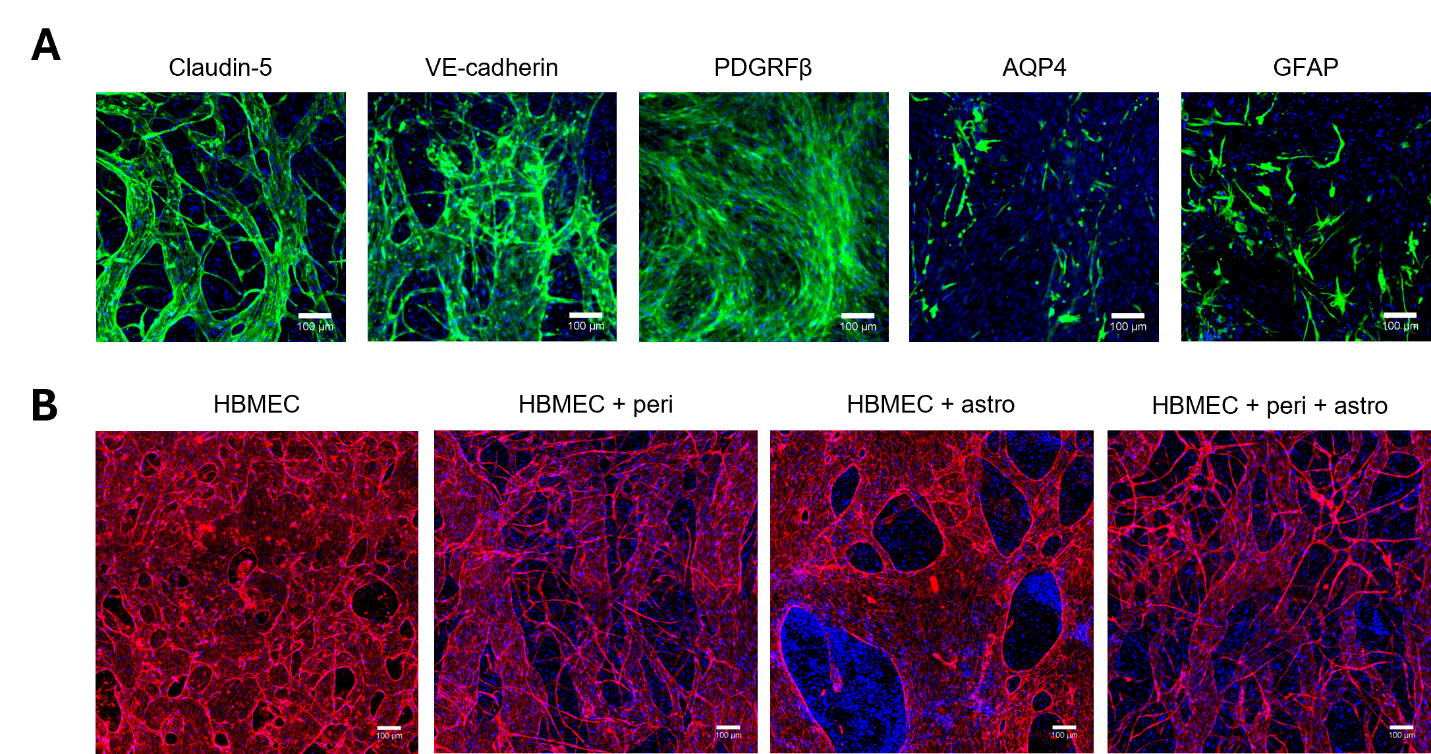

**Supplementary Figure 2 | Expression of endothelial, pericytic, and astrocytic markers. (A)** Co-cultures of HBMEC, astrocytes, and pericytes were fixed at day 14. HBMECs were stained for tight junction protein Claudin-5 and adherens junction protein VE-cadherin. Pericytes were stained for PDGFRβ and astrocytes were stained for AQ4 and GFAP. Scale bar = 100 µm. **(B)** Co-cultures of HBMEC, astrocytes, and pericytes were fixed at day 14 and stained for endothelial marker PECAM-1. Scale bar = 100 µm. PECAM-1 staining images were used for vascular network quantification shown in figure 2C-F.

**Supplementary video 1-4 | Unidirectional bead flow through BBB vascular networks.** Fluorescent beads (1-5 µm size) were perfused through the cerebral vascular networks. Bead flow through vascular networks consisting of (1) HBMEC, (2) HBMEC + pericytes, (3) HBMEC + astrocytes, and (4) HBMEC + astrocytes + pericytes was captured via high-speed fluorescent imaging.
